# PROTAC-Driven Protective Therapy increases the therapeutic window of anticancer drugs

**DOI:** 10.64898/2026.01.12.698947

**Authors:** Lucía Simón-Carrasco, Soledad Raya, Elena Pietrini, Manuel Luque-Perez, Marta del Rio Oliva, Iván V. Rosado, Andres J. Lopez-Contreras

## Abstract

Targeted protein degradation is emerging as a powerful anticancer therapy, mostly focused on eliminating oncogenic drivers. In contrast, we propose using PROTACs that exploit E3 ligase defects in cancer cells to selectively protect healthy tissues from the dose-limiting toxicity of anticancer drugs. We term this approach *PROTAC-Driven Protective Therapy* (PDPT). PDPT consists of a combinatorial treatment of a given anticancer compound with a PROTAC that promotes the degradation of proteins required for the drug-induced toxicity. Potential targets of protective PROTACs include drug uptake transporters, enzymes activating pro-drugs, and the actual drug target in cases that mediates the drug-induced toxicity. Notably, these protective PROTACs must be designed to recruit E3 ligases that are mutated or defective in the cancer cells while remain active in healthy tissues. As a proof of concept of our strategy, we used CRBN-recruiting and VHL-recruiting PROTACs to demonstrate that PARP1 degradation alleviates the cytotoxicity of PARP inhibitors (PARPi) in E3 ligase-proficient cells, while E3 ligase-deficient cancer cells remain fully sensitive. Remarkably, PDPT also protects primary human bone marrow progenitors from PARPi-induced toxicity, which are the most clinically relevant cells affected by PARPi-associated side effects in cancer patients, supporting the clinical relevance of this strategy. We further uncover that *TP53*-mutant cancers display critically low expression of the E3 ligase MDM2 and show inefficient MDM2-recruiting PROTAC activity. This tumor-intrinsic feature enables PDPT using MDM2-recruiting PROTACs in *TP53*-mutant cancers.

PDPT opens a new direction for targeted protein degradation by improving tolerability and expanding the therapeutic window of both established and future cancer therapies.

## Introduction

Personalized cancer therapy has emerged in the last decades as a promising approach to combat cancer. The general principle of personalized or targeted therapies is to exploit vulnerabilities of cancer cells, often provided by specific mutations, to selectively kill these cells over healthy cells. A well-known example of personalized cancer therapy is the use of Imatinib to selectively kill Chronic Myeloid Leukemia cells harboring the aberrant fusion tyrosine kinase BCR-ABL (ÓBrien et al. 2003, Druker et al. 1996). Among other relevant mutations, *TP53* is the most frequently mutated tumor suppressor gene in human cancers (Levine & Oren 2009), therefore, a specific cancer therapy targeting p53 deficiency could potentially benefit a large number of patients. However, despite vast research efforts and some notable examples targeting dysfunctional p53 for cancer treatment (Bykov et al. 2018, Peuget et al. 2024), efficient therapies selectively targeting cancers with *TP53* mutations are still lacking in the clinic.

PROTACs (proteolysis-targeting chimeras) are bivalent molecules that bind a target protein and an E3 ubiquitin ligase, aiming for the proteasomal degradation of the target protein. PROTACs and other protein degraders have emerged as a powerful tool for cancer therapy, enabling the selective degradation of previously “undruggable” proteins, including oncogenes with specific mutations such as KRAS^G12C^ (Bond et al. 2020) and KRAS^G12D^ (Zhou et al. 2024). Several PROTACs, such as the BET degrader dBET1 (Winter et al. 2015) or the BRD4 degrader ARV-825 (Wakita et al. 2020) have shown promising results in preclinical models. Moreover, some of them, such as the androgen receptor degrader ARV-110 (ClinicalTrials.gov ID NCT03888612, Snyder et al. 2025) and the estrogen receptor degrader ARV-471 (Phase III NCT05909397 and NCT05654623), are currently being tested in clinical trials.

Most of the efforts in this field have been directed towards the development of PROTACs targeting oncogenes or proteins required for cancer cells by recruiting E3 ligases, such as VHL, CRBN or MDM2, that are proficient in cancerous cells. However, applying this PROTAC technology to selectively target a tumor suppressor gene, such as *TP53*, which is often mutated or lost in cancer cells, is not straightforward. To circumvent this limitation, we have envisioned a novel approach: using PROTACs to selectively protect healthy tissues and thereby increase the therapeutic window of anticancer therapies. Undesired toxic effects of anticancer drugs on healthy tissues limit their efficacy by reducing the therapeutic window and often result in a reduction of the dose, duration, or even termination of the treatment. In this article, we describe the rationale of this novel approach that we name *PROTAC-Driven Protective Therapy (PDPT)* that exploits E3 ligase mutations or deficiencies in cancer cells, and we provide its proof of concept by combining PARP inhibitors (PARPi) with PARP1 degraders. Moreover, we uncover that MDM2-recruiting PROTACs are critically ineffective in *TP53*-mutant cancers, enabling the potential application of this strategy to a large number of cancer patients.

## Results

### Rationale of the PROTAC-Driven Protective Therapy (PDPT)

The PDPT consists of the combinatorial treatment of a known (*or novel*) anticancer drug with a PROTAC (Fig. 1). We propose that the precise, transient degradation of selected proteins in healthy tissues can mitigate undesired drug-induced toxicity. Critically, we introduce the concept of protective PROTACs, designed to degrade proteins that mediate drug-induced toxicity and to recruit an E3 ligase that is functional in healthy tissues but mutated or deficient in the tumor (Fig. 1A). As a result, the PROTAC promotes target degradation only in healthy cells, while sparing E3 ligase-deficient cancer cells, thereby increasing the therapeutic window of anticancer drugs. The majority of current PROTACs recruit the E3 ligases CRBN, VHL, or MDM2. Among these, VHL is frequently mutated in clear cell renal carcinoma, and notably we have found that MDM2 is deficient in *TP53*-mutant cancers.

**Figure 1.**
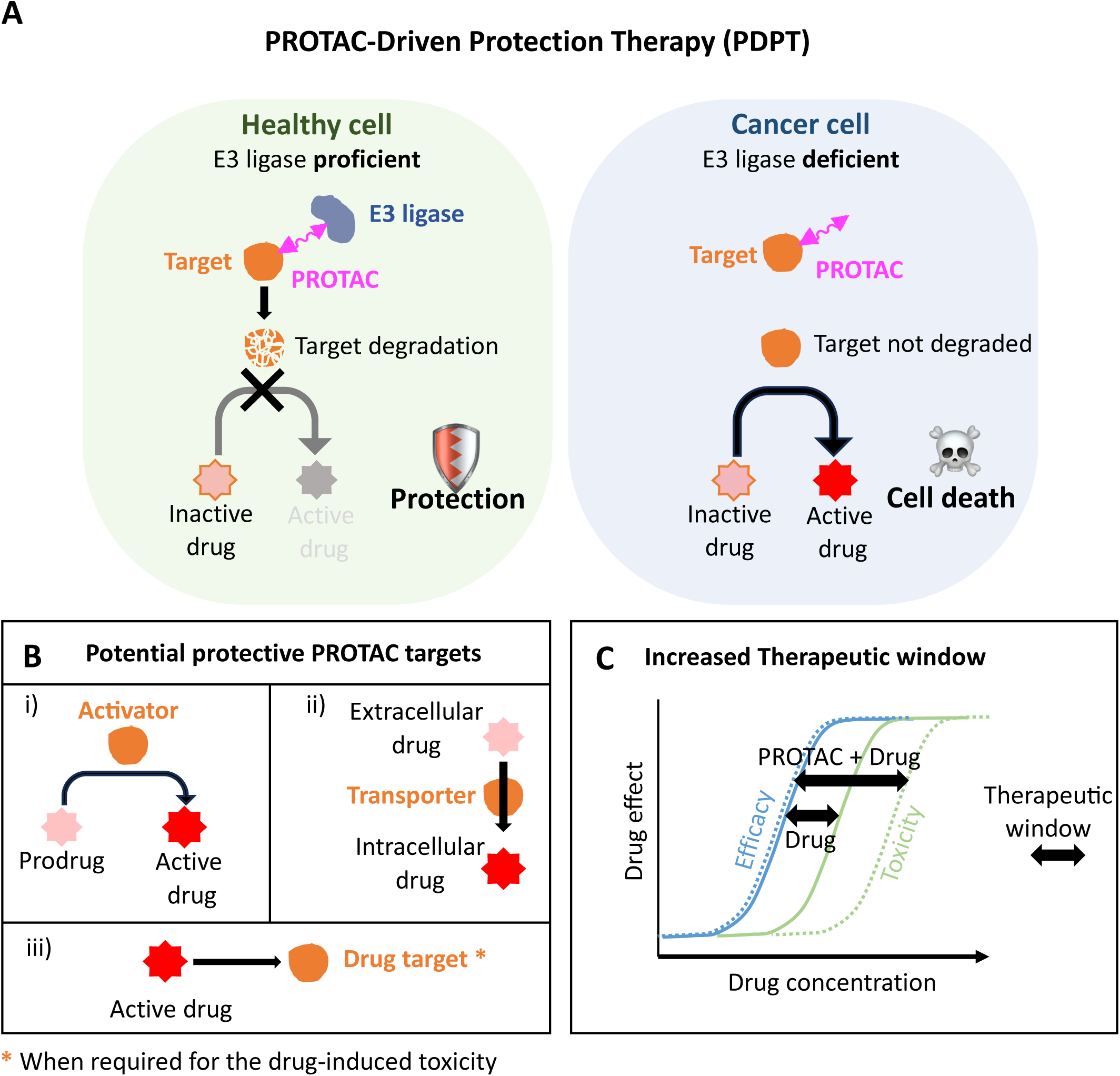
PROTAC-Driven Protective Therapy (PDPT) model. A) PDPT consists of a combinatorial treatment of an anticancer drug and a *protective* PROTAC. Healthy cells are E3 ligase proficient, and in the presence of the PROTAC, the target protein will be degraded. Loss of the target protein provides protection against the drug. In contrast, cancer cells that are deficient for the recruited E3 ligase cannot support PROTAC-mediated degradation of the target. Hence, the target protein will not be degraded, enabling the toxicity of the drug in cancer cells. B) Suitable protective PROTAC targets include: i) enzymes required to convert a prodrug into its active form, ii) transporters required to uptake the drug into the cell, and iii) direct drug targets that are required for the toxicity of the drug. C) Combined treatment with the PROTAC selectively reduces toxicity in healthy cells, thereby increasing the therapeutic window of anticancer drugs.

Target proteins suitable for PDPT include enzymes responsible for prodrug activation, transporters of the drugs, or the actual drug targets in cases where the protein target is responsible of the drug-induced toxicity (Fig. 1B). Importantly, the targeted protein must be non-essential and its loss should be well tolerated. Notably, the large number of CRISPR screen datasets available provides extensive information on genes whose loss confers drug resistance (https://orcs.thebiogrid.org) (Stark et al. 2006), which may facilitate the identification of potential PROTAC targets for this PDPT strategy.

We therefore propose that an appropriate therapeutic window can be achieved not only by increasing the sensitivity of cancer cells to certain drugs (traditional approaches) but also by increasing drug tolerance in healthy tissues, a conceptual shift enabled by the PDPT approach (Fig. 1C).

Two critical elements are required for PDPT: i) PROTAC-driven degradation of a protein that mediates drug toxicity in healthy tissues, and ii) E3 ligase defect in tumor cells to ensure selectivity. In the following sections, we demonstrate both principles experimentally.

### PARP inhibitors combined with PARP1 degraders as PDPT proof of concept

PARPi are used in the clinic for the treatment of breast cancer with *BRCA1/BRCA2* mutations and for the maintenance treatment of platinum-sensitive recurrent ovarian cancer (Cortesi et al. 2021,Tomao et al. 2019). In this context, patients undergoing prolonged PARPi therapy often present undesirable side effects such as anemia (LaFargue et al. 2019), due to PARPi-induced toxicity in hematopoietic progenitor cells (Maiorano et al. 2025). Interestingly, it has been reported that PARPi-induced toxicity is mainly mediated by PARP1-DNA trapping (Hopkins et al. 2019). In line with this, genome-wide CRISPR screens showed that PARP1 loss confers resistance to PARPi (Zimmermann et al. 2018, BioGRID ORCS Screen ID: 1352). Thus, it would be clinically relevant to increase PARPi tolerance in healthy tissues, increasing their therapeutic window. For the proof of concept of the PDPT strategy, first, we aimed to demonstrate that targeting the appropriate protein with a PROTAC can be used to protect cells from the toxicity of an anticancer drug. We used a CRBN-recruiting PARP1 degrader, the SK-575 PROTAC (Cao et al. 2020) (Fig. 2A), to alleviate PARPi-induced toxicity. The SK-575 PROTAC is formed by Olaparib (that acts as PARP1 binder) fused through a linker to Thalidomide (CRBN binder) and has shown high *in vitro* and *in vivo* efficacy and selectivity in degrading PARP1 (Cao et al. 2020). Importantly for our strategy, several studies indicate that PARP1 depletion is well tolerated *in vivo*, both genetically (Wang et al. 1995, Farrés et al. 2015) and using PROTACs (Li et al. 2022).

**Figure 2.**
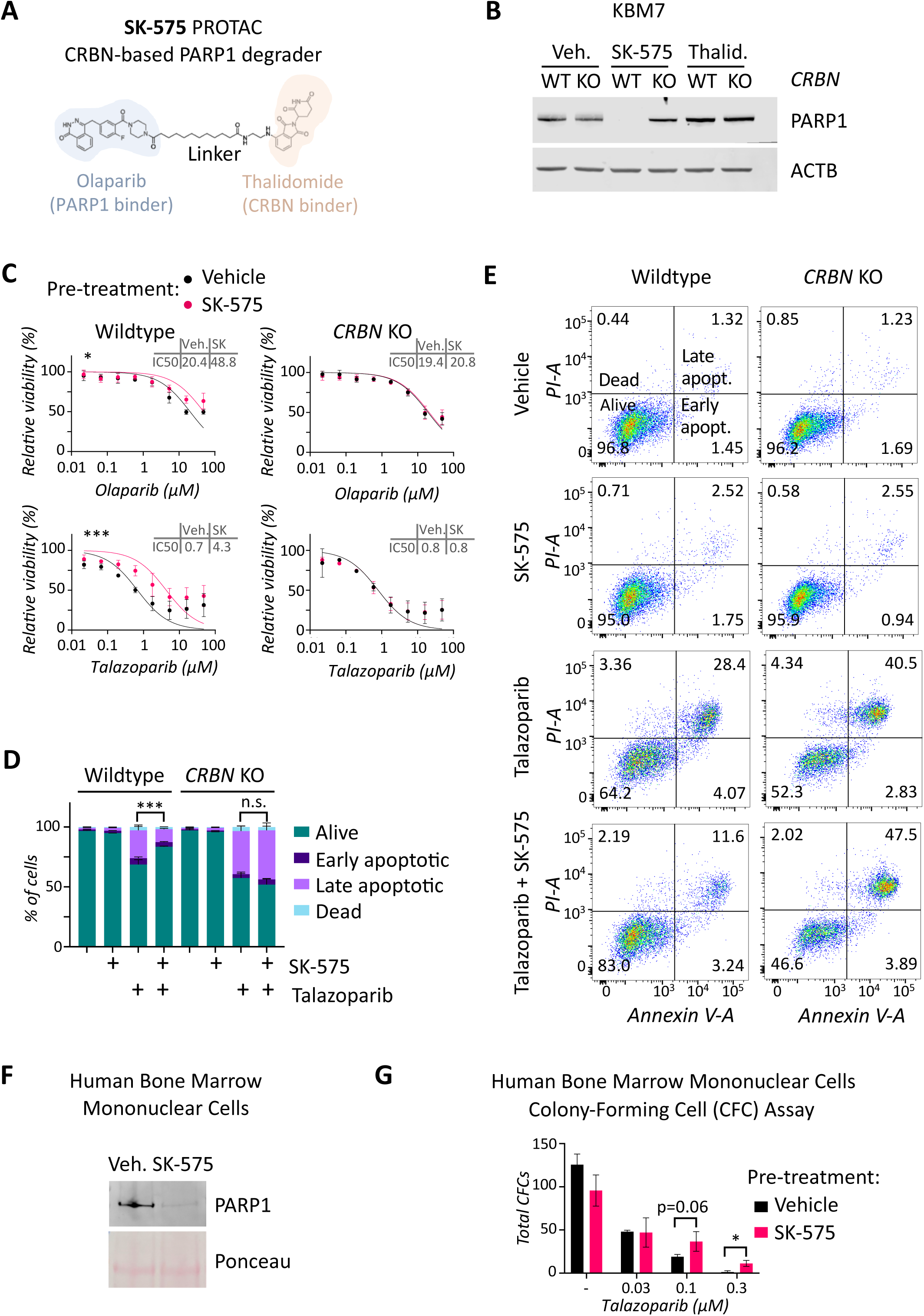
PARP1 degradation by a CRBN-recruiting PROTAC alleviates PARPi-induced cell toxicity. A) Structure of SK-575 PROTAC formed by the PARPi Olaparib linked to Thalidomide, a CRBN binder. B) Western blot analysis of PARP1 expression in isogenic *CRBN* WT and KO KBM7 cells treated with 100 nM SK-575 or Thalidomide for 48h. ACTB was used as loading control. C) Viability assay by MTT performed in *CRBN* WT or KO KBM7 cells pre-treated with 100 nM SK-575 for 24 h followed by a treatment with both SK-575 and the indicated doses of Olaparib or Talazoparib for another 24 h. Data are normalized to vehicle or SK-575 pre-treated cells and correspond to three biological replicates. IC50s were calculated with non-linear regression analysis and are shown in the figure (µM). Statistical significance was determined with Extra sum-of-squares F test. D) Analysis of percentage of viability as assessed by Annexin V/PI staining using flow cytometry analysis from three biological replicates. Statistical significance was determined with Ordinary one-way ANOVA and Tukey’s multiple comparisons test. E) Representative images of the Annexin V/PI staining. Percentage of cells for each population are shown. F) Western blot showing PARP1 degradation in human bone marrow mononuclear cells treated with 100 nM SK-575 for 48 h. G) Colony-forming cells (CFC) assay of human bone marrow mononuclear cells pre-treated with 100 nM SK-575 24 h before seeding them in MethoCult semi-solid methylcellulose-based media with the indicated doses of Talazoparib with or without 100 nM SK-575. Colonies were quantified after 14 days. Error bars indicate mean ± s.d. (n = 3). Statistical significance was determined with unpaired t-tests. In all the figures * = *p* < 0.05, and *** = *p* < 0.001.

We used KBM7 *CRBN* WT and KO isogenic cells to compare the combined effect of SK-575 and two different PARPi used in the clinic (Olaparib and Talazoparib) in E3 ligase proficient and deficient cells. The SK-575 PROTAC at 100 nM induced complete PARP1 degradation in KBM7 parental cells within 48 h, while it did not affect PARP1 levels in the KBM7 *CRBN* KO cells (Fig. 2B). Thalidomide alone at 100 nM did not affect PARP1 levels (Fig. 2B).

To ensure efficient PARP1 degradation prior to PARPi exposure, cells were pre-treated with SK-575 at 100 nM for 24 h before exposure to the indicated PARPi doses in combination with SK-575 for an additional 24 h, and cell viability was determined by MTT assay. Notably, SK-575 reduced the PARPi-induced toxicity in *CRBN* WT cells, while having no effect in *CRBN* KO cells (Fig. 2C), indicating that PARP1 degradation is responsible of the protection. In *CRBN* WT cells, SK-575 increased Olaparib IC50 from 20.37 to 48.83 μM, and Talazoparib from 0.68 to 4.32 μM (Fig. 2C). The SK-575 PROTAC provided stronger protection from Talazoparib-induced toxicity likely due to its reported higher PARP1 trapping activity compared to Olaparib (Hopkins et al. 2015). Apoptosis analysis by FACS using Annexin-V/PI staining corroborated the PROTAC protective effect against Talazoparib-induced toxicity selectively in *CRBN* WT cells (Figs. 2D and 2E). Similar findings were obtained using a VHL-recruiting PARP1 degrader 180055 (Chen et al. 2024) (Fig. S1A) in *VHL* WT and KO KBM7 cell lines (Fig. S1B). The PARP1 degrader alleviated Talazoparib-induced toxicity selectively in *VHL* WT cells (Fig. S1C). Interestingly, when the pretreatment with SK-575 was not included, and SK-575 and Talazoparib were added simultaneously, the protective effect was not observed (Fig. S2A), suggesting that SK-575 and Talazoparib may compete for PARP1 binding.

These results demonstrate that PDPT can protect E3 ligase proficient cells from a given chemotherapy without reducing its efficacy in E3 ligase mutant cells.

### PROTAC-driven PARP1 depletion protects human hematopoietic progenitor cells from PARPi

The study of *CRBN* (and *VHL*) WT and KO isogenic cell lines allowed us to validate the selectivity of the PDPT strategy towards specific E3 ligase deficiencies. However, for PARPi, the most clinically relevant undesirable toxicity occurs in hematopoietic progenitor cells (Maiorano et al. 2025). Therefore, we sought to evaluate whether the SK-575 PARP1 degrader could provide protection in this cellular context. To do this, we performed colony-forming cells (CFC) assays using human bone marrow mononuclear cells. Cells were pretreated (or not) with SK-575 at 100 nM for 24 h, and then subjected to the indicated doses of Talazoparib, either alone or in combination with SK-575, in a semi-solid matrix that supports clonal progenitor growth. Colonies were quantified after 14 days in culture. The treatment with SK-575 at 100 nM achieved a robust PARP1 degradation (Fig. 2F). As expected, hematopoietic progenitor cells displayed high sensitivity to Talazoparib and, remarkably, the PARP1 degrader attenuated Talazoparib-induced toxicity at the higher concentrations tested (Fig. 2G). In line with our results in KBM7 cells, when the cells were not pretreated with SK-575, the SK-575-mediated protection from Talazoparib was attenuated (Fig. S2B).

These results indicate that PDPT could protect clinically relevant healthy tissues from chemotherapy-induced toxicity. We next aimed to identify cancer-specific E3 ligase defects that could be exploited to expand PDPT to a large number of cancer patients.

### Loss of p53 impairs the activity of MDM2-recruiting PROTACs

A second prerequisite for PDPT is the existence of E3 ligases that are specifically defective in tumors but functional in healthy tissues. In the previous sections we have used CRBN and VHL-recruiting PROTACs. The potential clinical application of the PDPT using CRBN-recruiting PROTACs is limited by the low number of patients with *CRBN* mutations, although such alterations have been described for multiple myeloma patients resistant to immunomodulatory therapies with Thalidomide, Lenalidomide, and other compounds targeting CRBN (Gooding et al. 2021). On the other hand, *VHL* is frequently mutated in clear cell renal carcinoma (Cancer Genome Atlas Research Network, 2013), opening the possibility for PDPT based on VHL-recruiting PROTACs in this patient group. Moreover, given that MDM2, which is another E3 ligase currently used in PROTAC technology, is transcriptionally regulated by p53 (Barak et al., 1993), we hypothesized that *TP53*-mutant cancers may display reduced MDM2 levels and therefore impaired MDM2-recruiting PROTAC activity. Such MDM2 deficiency would open a therapeutic opportunity for treating *TP53*-mutant cancers with PDPT strategy, enabling the potential application of PDPT to a large proportion of cancer patients.

To experimentally validate this hypothesis, we used HCT116 *TP53* WT and KO isogenic cell lines, and an available MDM2-recruiting BRD4 PROTAC A1874, formed by the MDM2 antagonist Idasanutlin joined through a linker to the BRD4 inhibitor JQ1 (Hines et al. 2019) (Fig. 3A). Note that BRD4 is not a relevant target for PDPT strategies and we used this PROTAC only to characterize the activity of MDM2-recruiting PROTACs in the context of p53 deficiency. We found that A1874 induced BRD4 degradation in HCT116 parental cells (*TP53* WT) while it was ineffective in the *TP53* KO cells (Fig. 3B). This differential effect correlated with lower MDM2 expression in the *TP53* KO cells at the protein and mRNA levels (Figs. 3B, 3C and 3D). Moreover, the A1874 PROTAC markedly induced MDM2 expression selectively in the *TP53* WT cells (Figs. 3B, 3C and 3D), further increasing the differential levels of MDM2 between *TP53* WT and KO HCT116 cells. The MDM2 induction by the A1874 PROTAC is likely caused by p53 stabilization mediated by the Idasanutlin component of the molecule (Ding et al. 2013). Indeed, A1874 induced p53 stabilization and p21 induction in *TP53* WT cells. Intriguingly, it also caused a modest increase in p21 levels in *TP53* KO cells (Fig. 3B), probably induced by p73 stabilization as previously described (Ongkeko et al. 1999, Lau et al. 2008).

**Figure 3.**
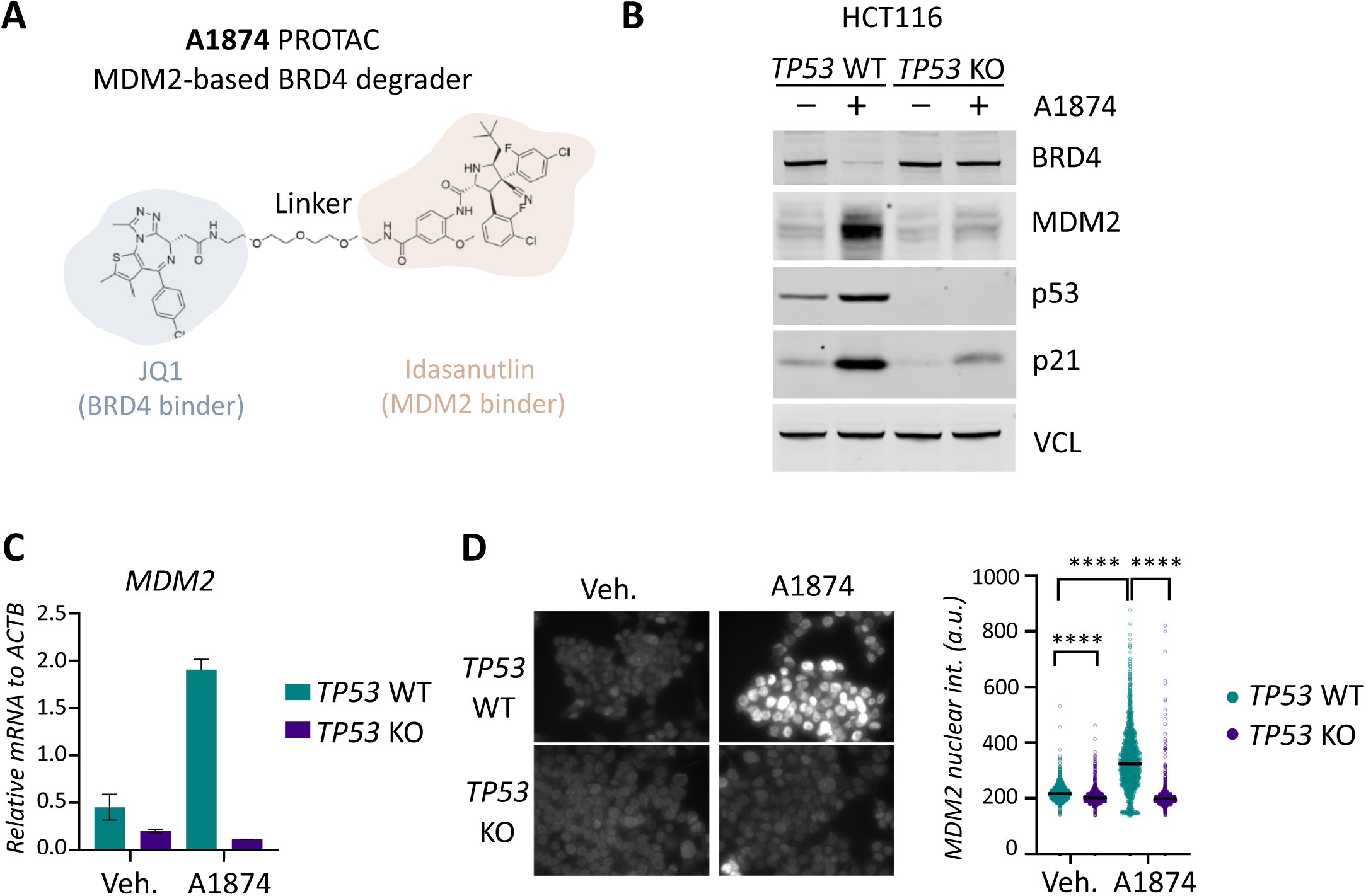
p53 loss impairs MDM2-recruiting PROTACs activity. A) Structure of the MDM2-recruiting BRD4 degrader A1874 formed by JQ1 (a BRD4 inhibitor) and Idasanutlin (an MDM2 antagonist). B) BRD4, MDM2, p53 and p21 protein levels in isogenic *TP53* WT and KO HCT116 cells treated with 1 µM A1874 for 24 h, assessed by western blot. Vinculin (VLC) was used as loading control. C) *MDM2* mRNA expression levels in *TP53* WT and *TP53* KO HCT116 cells treated with 1 µM A1874 for 24 h assessed by qPCR. Error bars indicate mean ± s.d. (n = 3). D) Representative immunofluorescence images of MDM2 levels in *TP53* WT and KO HCT116 cells upon treatment with vehicle or 1 µM A1874 for 24 h. Quantification of MDM2 nuclear intensity from 1600 cells is shown on the right. Statistical significance was determined with unpaired t-tests. **** = *p* < 0.0001.

Given these observations, we next investigated whether the effect was restricted to complete p53 loss or also extended to *TP53* missense mutations frequently found in human cancers. To this end, we transduced *TP53* KO HCT116 cells with lentiviral plasmids expressing p53 WT or the p53 mutant variants R175H, R249S, R273H, and R280K. The expression of p53 mutant versions caused a robust accumulation of p53 (Fig. S3A). Remarkably, the MDM2-recruiting PROTAC A1874 was still inefficient in degrading BRD4, consistent with the absence of MDM2 activation (Fig. S3B). Although we did not observe a rescue of PROTAC activity with the p53 WT-expressing plasmid, this was likely due to the fact that p53-WT overexpression severely impaired cell proliferation, in contrast to p53-mutant overexpression. Overall, these data suggest that MDM2-recruiting PROTACs are inefficient in cancer cells harboring *TP53* missense mutations.

### MDM2-recruiting PROTACs are defective in cancer cells harboring *TP53* mutations

To evaluate the generality of our observations, we next sought to corroborate our findings in human cancer samples and in *TP53*-mutant human cancer cell lines derived from different tissues. Analyses of the TCGA pan-cancer dataset showed that *TP53*-mutant cancers globally display reduced mRNA levels of *MDM2* (Fig. S4A and S4B). This correlation was also observed in individual TCGA datasets from colorectal, breast cancer and melanoma tumors (Fig. S4A and S4B). Given the clinical relevance of PARPi in ovarian cancer, we aimed to analyze whether this association was also present in TCGA ovarian cancer dataset. However, this analysis could not be done due to the limited availability of *MDM2* expression data in *TP53*-mutant ovarian tumors in this dataset (Fig. S4A).

For this reason, and considering the relevance of ovarian cancer models to PARPi therapy, we complemented these cancer database analyses by experimentally examining a panel of colorectal and ovarian cancer cell lines harboring -or not-*TP53* mutations. Consistent with our *in-silico* observations, *TP53*-mutant cell lines showed impaired responses to MDM2-recruiting PROTACs. The BRD4 degrader A1874 was ineffective in all the *TP53*-mutant cell lines tested (Fig. 4A). In contrast, this PROTAC induced robust BRD4 degradation in *TP53*-WT cell lines, including the colorectal cancer cell lines HCT116 and RKO, the ovarian cancer cell line A2780, and the primary cell lines BJ and RPE1 (Fig. 4A). Notably, *MDM2* mRNA levels were lower in the *TP53*-mutant colon cancer cell lines (DLD1, HT29) compared to the *TP53*-WT cell lines (HCT116 and RKO), and this differential MDM2 expression was exacerbated upon A1874 treatment (Fig. 4B), likely due to the Idasanutlin moiety of the PROTAC. Similar differences in *MDM2* expression were observed when comparing *TP53*-WT (A2780) and *TP53*-mutant (ES2, OVCAR8, SKOV3) ovarian cancer cell lines (Fig. 4B). Remarkably, the MDM2-recruiting PROTAC induced *MDM2* mRNA expression in all the *TP53*-WT cell lines (HCT116, RKO, A2780, BJ and RPE1), but in none of the *TP53*-mutant cell lines tested (Fig. 4B).

**Figure 4.**
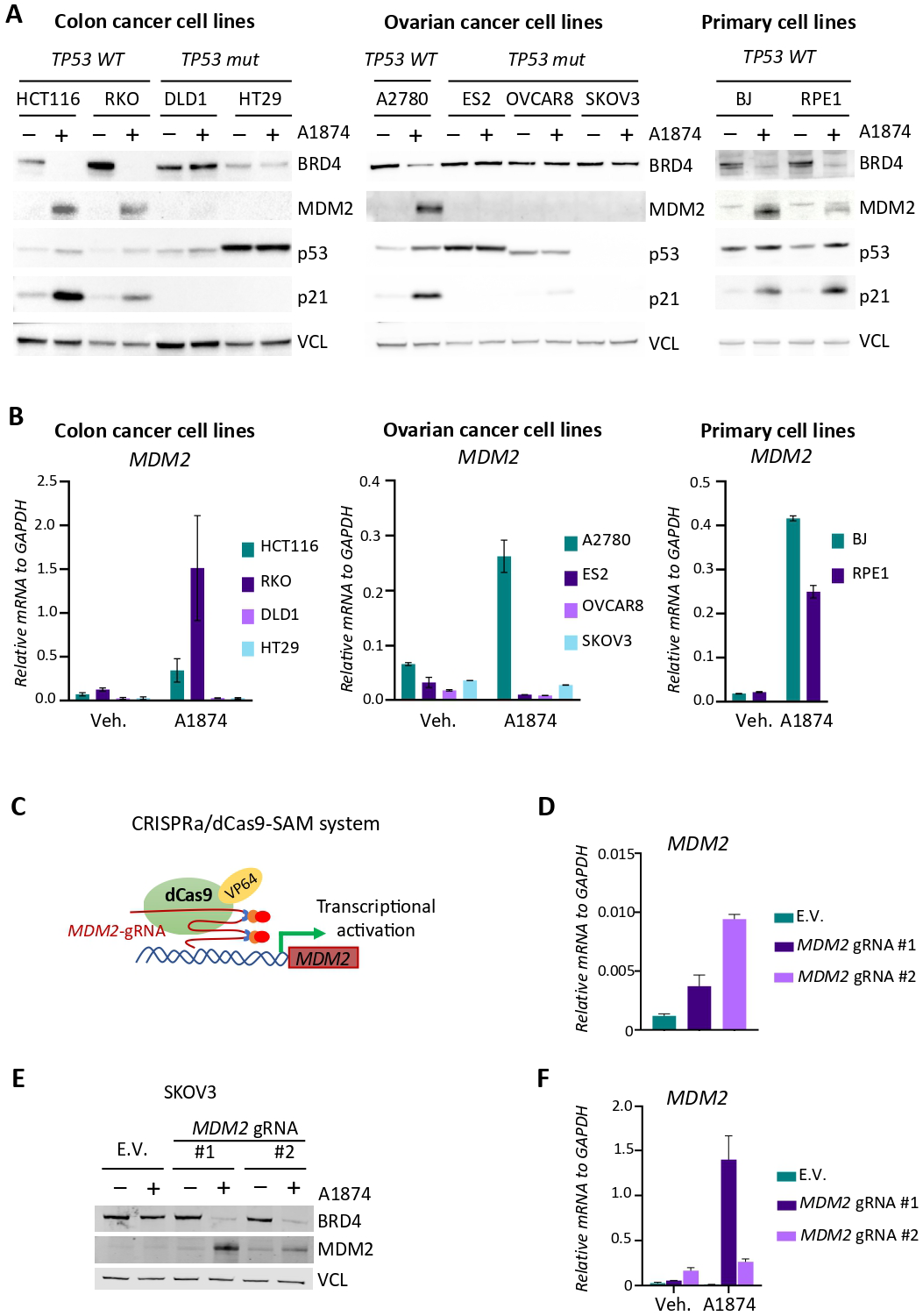
MDM2-recruiting PROTACs are ineffective in *TP53*-mutant cancer cell lines. A) BRD4, MDM2, p53, and p21 protein levels in a panel of cancer cell lines treated with vehicle or 1 µM A1874 for 24 h, assessed by western blot. Vinculin was used as loading control. Left panel displays colon cancer lines, including *TP53* WT (HCT116 and RKO) and mutant (DLD1 and HT29) cells. Middle panel displays ovarian cancer cell lines, including *TP53* WT (A2780) and mutant (ES2, OVCAR8 and SKOV3) cells. Right panel displays BJ and RPE1 p53-proficient primary cell lines. B) *MDM2* mRNA expression levels in the indicated cell lines treated with vehicle or 1 µM A1874 for 24 h, assessed by qPCR. Error bars indicate mean ± s.d. (n = 3). C) Schematic representation of the CRISPRa/dCas9-SAM system. It consists of a dead Cas9 (dCas9) fused to the transcriptional activator VP64. The gRNA targeting MDM2 promoter forms a complex with MS2 (blue) and recruits transcriptional p65 (orange) and HSF1 (red) activators. This induces *MDM2* expression from the endogenous locus. D) *MDM2* mRNA expression levels in SKOV3 transduced with E.V. or gRNAs targeting *MDM2* promoter, assessed by qPCR. Error bars indicate mean ± s.d. (n = 3). E) BRD4, and MDM2 protein levels in empty vector (E.V.)-transduced or MDM2-overexpressing SKOV3 cell lines treated with 1 µM A1874 for 24 h, assessed by western blot. Vinculin was used as loading control. F) *MDM2* mRNA expression levels in the indicated cells treated with vehicle or 1 µM A1874 for 24 h, assessed by qPCR. Error bars indicate mean ± s.d. (n = 3).

Overall, the strong correlation between *TP53* mutations, *MDM2* expression, and MDM2-recruiting PROTAC activity indicates that MDM2 deficiency is a specific tumor-intrinsic feature of *TP53*-mutant cancers that can be exploited for PDPT.

### MDM2 restoration rescues PROTAC activity in *TP53*-mutant cancers

The correlation observed between MDM2 levels and the efficiency of the MDM2-recruiting BRD4 PROTAC across multiple cell lines suggested that reduced *MDM2* expression is a key determinant impairing PROTAC activity in p53-deficient cells. However, given that Idasanutlin binds the interface between MDM2 and p53, we wondered whether restoring MDM2 expression alone was sufficient to rescue MDM2-recruiting PROTAC activity in a p53-deficient context. To this end, we used the dCas9-SAM activation system to upregulate *MDM2* expression in the *TP53*-mutant SKOV3 ovarian cancer cell line. SKOV3 harbors a *TP53* early truncating mutation (c.267delC, p.S90fsX33) (Ikediobi et al. 2006), resulting in the complete loss of functional p53 protein, as shown in Fig. 4A. We transduced SKOV3 cells with the dCas9-SAM system and either an empty vector (E.V.) or vectors encoding two gRNAs targeting the *MDM2* promoter (Fig. 4C). Both gRNAs induced *MDM2* expression under basal conditions (Fig. 4D), and this effect was further amplified upon A1874 treatment (Figs. 4E and 4F). Notably, the MDM2 induction achieved by both gRNAs was sufficient to promote efficient BRD4 degradation by the A1874 PROTAC molecule (Fig. 4E). These data confirm that insufficient MDM2 expression is the critical molecular event impairing MDM2-recruiting PROTAC activity in p53-deficient cells.

## Discussion

In this study, we introduce a conceptually novel therapeutic strategy, PROTAC-Driven Protective Therapy (PDPT) that proposes the use of targeted protein degradation not to sensitize cancer cells, but to selectively protect healthy tissues, thereby increasing the therapeutic window of anticancer drugs. Our results demonstrate the feasibility of this approach and show how E3 ligase mutations or deficiencies in cancer cells enable selective protein degradation in non-tumoral cells.

To our knowledge, this strategy entails a paradigmatic shift in the current rationale of protein degradation-based strategies for cancer therapy. Most PROTACs developed to date aim to eliminate oncogenes or pro-proliferative factors in cancer cells, and are designed to recruit E3 ligases that are active in tumors. On the contrary, PDPT exploits E3 ligases that are mutated or inactive in cancer, thereby restricting the target degradation to healthy tissues. Notably, several E3 ligases are mutated in cancer, for instance VHL in clear cell renal carcinoma, FBXW7 in endometrial, colorectal and cervical cancers, or BRCA1 in breast and ovarian cancers. Of these three E3 ligases, only VHL-recruiting PROTACs have been designed to date. Moreover, as we demonstrate here, MDM2 is defective in *TP53*-mutant cancers, enabling the use of MDM2-recruiting PROTACs for PDPT. Further, it would be beneficial for PDPT strategies to develop novel PROTACs recruiting other E3 ligases frequently mutated in cancer such as FBXW7 or BRCA1.

For PDPT to be effective, the targeted protein must contribute to the drug-induced toxicity and its transient depletion must not cause further toxic effects in healthy tissues. PARP1 fulfills these criteria. Genetic studies indicate that PARP1 loss is well tolerated *in vivo* (Wang et al. 1995, Farrés et al. 2015), and PARPi toxicity is mediated by PARP1-DNA trapping (Hopkins et al. 2019). Using a CRBN-recruiting PARP1 degrader, we demonstrated that PARP1 loss markedly reduces PARPi-induced toxicity in cell lines and, critically, in human hematopoietic progenitor cells, which are the most relevant cells suffering PARPi-toxicity in cancer patients with prolonged PARPi therapy (Maiorano et al. 2025). We also recapitulated the protection with a VHL-recruiting PROTAC, showing that this effect is not dependent on the E3 ligase. Interestingly, we found that the protective effect relies on the appropriate PARP1 degradation prior to PARPi administration, indicating the need for careful schedule design in future *in vivo* PDPT studies. However, these timing considerations might not be required in those cases where the PROTAC and the drug target different proteins.

Notably, PDPT is not limited to PARP1 and PARPi. Indeed, we believe that PDPT has the potential to be scalable to a large number of protein targets and several E3 ligases. Other suitable protein targets are enzymes required for prodrug activation, transporters required for the cellular drug uptake, and other factors mediating the toxicity of a given anticancer drug, including the actual drug target in some cases (as in the example of PARP1 and PARPi). One potential application of this strategy would be the development of a protective PROTAC targeting deoxycytidine kinase (dCK) to use in combination with Gemcitabine in PDPT. Given that dCK is required to convert Gemcitabine into its active form (Toschi et al. 2005), and its loss confers resistance to this anticancer compound (Lau et al. 2020, BioGRID ORCS Screen ID: 2379), dCK selective degradation in healthy tissues could protect them from Gemcitabine toxicity while tumor cells remained sensitive. Of note, dKC loss or pharmacological inhibition are well tolerated *in vivo* (Toy et al. 2010, Poddar et al. 2020), supporting the use of future dCK degraders for PDPT strategies.

As mentioned above, the E3 ligase used in protective PROTACs must be mutated or defective in the cancer cells and, critically, must be expressed in the tissues where the anticancer compounds cause the major off-target toxicities. In this regard, relevant E3 ligases for the PDPT strategy including VHL, MDM2, BRCA1, and FBXW7 are constitutively expressed in all human tissues (data from https://www.proteinatlas.org/, (Uhlén et al. 2015)), supporting their use in the design of protective PROTACs.

A key mechanistic insight from our work is the characterization that p53-deficient cancers exhibit a profound defect in MDM2-recruiting PROTAC activity, explained by critically reduced MDM2 levels. Indeed, we found this deficiency across multiple *TP53* mutations and cancer types. Importantly, this intrinsic and mutation-driven feature of *TP53*-mutant cells enables the potential use of MDM2-based PDPT strategies in a large number of cancer patients with *TP53* mutations. In addition, this finding highlights that *TP53*-mutant cancers may be refractory to conventional protein targeted strategies with MDM2-recruiting PROTACs.

Another aspect to take into consideration for the development of protective PROTACs is the design of E3 ligase binders. Particularly, current MDM2-recruiting PROTACs rely on Nutlin or Idasanutlin derivatives, which cause p53 stabilization and MDM2 overexpression. We have seen that this effect may be positive to further upregulate MDM2 in healthy tissues increasing the MDM2-recruiting PROTAC activity. However, p53 stabilization may also lead to dose-dependent toxicities (Konopleva et al. 2020). Therefore, future PDPT strategies using MDM2-recruiting PROTACs will require identifying PROTACs that efficiently degrade their targets at concentrations that do not significantly induce p53-mediated toxicity. Additionally, it would be relevant for future PDPT strategies to develop different MDM2 binders that do not activate p53 at all. Despite this potential limitation, it should be noted that Idasanutlin by itself has been used in several clinical trials (Konopleva et al. 2020) and PROTACs are usually efficient at lower doses than traditional drugs due to their catalytic mode of action.

Interestingly, PDPT could also serve as a second line of treatment for tumors that acquire resistance to traditional PROTAC-based cancer therapies caused by E3 ligase loss or mutation. The growing field developing PROTACs to target oncoproteins in tumor cells, will likely translate into the emergence of treatment-derived mutations in E3 ligases in resistant cancer cells, similarly to the *CRBN* mutations occurring in multiple myeloma patients treated with immunomodulators targeting CRBN (Gooding et al. 2021). The appearance of E3 ligase mutations in resistant cancer cells may enable the shift to a PDPT strategy targeting the same E3 ligase to protect healthy tissues.

We would like to emphasize that our PDPT model offers opportunities to selectively reduce the undesirable toxic effects of certain anticancer drugs currently used in clinic, potentially increasing their therapeutic window and applications. Additionally, PDPT may allow the development of novel otherwise cytotoxic compounds into clinically functional targeted therapies. This perspective may expand the possibilities for drug development, especially in early stages where toxicities frequently limit the progression of novel candidates.

In conclusion, our study introduces a conceptually novel approach using targeted protein degradation that exploits tumor-specific E3 ligase deficiencies to improve the tolerability of cancer therapies. We anticipate that the principles described here will stimulate future studies to develop protective PROTACs with a renewed view that would shift and expand the relevant E3 ligases and proteins targeted for cancer therapy purposes.

## Methods

### Cell lines and cell culture

Human Myeloid Leukemia KBM7 parental and the isogenic *CRBN* KO and *VHL* KO cell lines were kindly provided by Dr. Cristina Mayor-Ruiz (IRB Barcelona) (Mayor-Ruiz et al. 2019). Human colorectal HCT116 parental and isogenic *TP53* KO cell lines were kindly provided by the Vogelstein laboratory (Bunz et al. 1998). All cell lines tested negative for *Mycoplasma* contamination. HCT116, RKO and BJ were grown in DMEM containing 2mM glutamine, 10% FBS, and 1% penicillin/streptomycin, DLD1, A2780 and OVCAR8 were grown in RPMI containing 2mM glutamine, 10% FBS, and 1% penicillin/streptomycin, HT29, ES2 and SKOV3 were grown in McCoy’s 5A containing 1.5mM glutamine, 10% FBS, and 1% penicillin/streptomycin, KBM7 was grown in IMDM containing 4mM glutamine, 10% FBS, and 1% penicillin/streptomycin and RPE1 was grown in DMEM/F12 containing 2mM glutamine, 10% FBS, and 1% penicillin/streptomycin.

### Western blot

Cells were lysed using modified RIPA lysis buffer (50mM Tris-Cl pH 7.5, 150 mM NaCl, 1 mM EDTA pH8, 1% NP-40, and 0.1% sodium deoxycholate) supplemented with Roche complete protease inhibitor cocktail (Roche, 04693116001), 5mM sodium fluoride (SIGMA, 201154), 5mM β-glycerophosphate disodium (SIGMA, G5422) and 2mM sodium orthovanadate (SIGMA, S6508). Protein concentrations were determined using DC Protein Assay reagents (BIORAD) with BSA as a protein standard. Equal amounts of protein were resolved on NuPAGE™ Bis-Tris Mini Protein Gels, 4–12% (Invitrogen) and transferred onto nitrocellulose membranes (Amersham Protran 0.2 µm NC). Membranes were blocked with 5% milk in PBST and incubated with the indicated primary antibodies overnight. The next day, they were incubated with horseradish peroxidase–conjugated secondary antibodies or fluorescently labelled secondary antibodies, and the signal was detected by chemiluminescence using ImageQuant 800 (AMERSHAM) or fluorescence in Odyssey CLx (LI-COR), respectively. Primary antibodies used were: BRD4 (A301-985A100 Bethyl 1:1000), MDM2 (SMP14) (sc-965 Santa Cruz 1:1000), p53 (DO-1) (sc-126 Santa Cruz 1:1000), p21 Waf1/Cip1 (12D1) Rabbit Monoclonal Antibody (2947 Cell Signaling 1:1000), Vinculin (7F9) (sc-73614 Santa Cruz 1:10000) and PARP-1 (B-10) (sc-74470 Santa Cruz 1:1000).

### MTT viability assay

Cells were seeded in 96-well plates at a density of 1×10^4^ cells per well. In each experiment, three technical replicates were seeded for each condition. For the 3-(4,5dimethylthiazol-2-yl)-2,5-diphenyltetrazoliumbromide (MTT) assay, cells were seeded in 96-well plates at a density of 5×10^4^ cells per well. 48 h later, cells were incubated with MTT solution (Sigma-Aldrich, 0.5mg/ml) for 75 minutes. Formazan was solubilized in 8.3 µM HCl/ 10% SDS, and absorbances at 570 nm and at 690 nm (background) were measured using a Varioskan Lux plate reader (Thermo Fisher Scientific).

### Drug treatment of cell lines

Cells were treated with the following compounds at the indicated doses: BRD4 PROTAC A1874 (HY-114305, MedChemExpress) at 1 µM; PARP1 PROTACs SK-575 (HY-139156, MedChemExpress) at 100 nM and 180055 (HY-170620, MedChemExpress) at 1 µM; and PARP inhibitors Talazoparib (HY-16106, MedChemExpress) and Olaparib (HY-10162, MedChemExpress) at concentrations ranging from 48 µM to 0.02 µM in 3-fold dilutions. Equivalent concentration of DMSO were used as vehicle controls. Normalized absorbance values relative to vehicle-only or PROTAC-only treated cells were plotted using GraphPad Prism.

### Annexin V/PI apoptosis assay

Apoptosis was evaluated by flow cytometry analysis using an Annexin V-FITC apoptosis detection kit from Immunostep S.L. (ANXVKF) following the manufacturer’s instructions. Briefly, 5×10^5^ cells were resuspended in binding buffer, stained with Annexin V-FITC and Propidium Iodide for 15 min in the dark, and washed with binding buffer. Cells were analyzed using a LSRFortessa Cell Analyzer cytometer (BD Biosciences). Data were analyzed using FlowJo software.

### Colony-Forming Cells assay

Colony-Forming Cells (CFC) assays were conducted at StemCell Technologies. Frozen human bone marrow mononuclear cells (MNCs) were thawed, washed, counted, and assessed for viability. Cells were pretreated (or not) for 24h with 100nM SK-575. Next, an appropriate number of MNCs were added to each aliquot of MethoCult™ media (GF H84434, Stemcell Technologies) containing Talazoparib or vehicle, with or without SK-575. The contents of each tube of MethoCult™ media were vortexed before being plated into three replicate wells of a 6-well STEMvision™ SmartDish™, which was placed at 37°C, 5% CO2 for 14 days. After the appropriate incubation period, the total number of erythroid (CFU-E and BFU-E), myeloid (CFU-GM) and, multi-potential progenitor (CFU-GEMM) colonies was evaluated by trained personnel and enumerated based on morphology.

### qRT-PCR

Total RNA was extracted using the RNeasy Mini Kit (Qiagen) and reverse-transcribed using the Maxima First Strand cDNA synthesis kit (K1641, Thermo Scientific) following the manufacturer’s instructions. Quantitative real-time PCR reactions were performed on a 7500 Fast real-time PCR system (Applied Biosystems) using the SYBR Green PCR Master Mix (Applied Biosystems). Values were quantified according to the ΔΔCt method, and *ACTB* or *GAPDH* was used for normalization. The primer sequences used are: human *MDM2* forward 5’-CTGTGTGTAATAAGGGAGATATGTTGTG-3’ and reverse 5’-GAATGTTCACTTACACCAGCATCAA-3’, human *GAPDH* forward 5’-TGCACCACCAACTGCTTAGC-3’ and reverse 5’-GGCATGGACTGTGGTCATGAG-3’, human *ACTB* forward 5’-CTGGAACGGTGAAGGTGACA-3’and reverse 5’-AAGGGACTTCCTGTAACAATGCA-3’.

### Immunofluorescence

Cells were seeded in polystyrene 96-well microplate μClear® (Greiner, 655090) 24 h prior to treatment. They were then treated with 1 µM A1874 PROTAC. After 24 h, cells were fixed with 4% PFA in PBS for 10 minutes at RT, permeabilized with 0.5% Triton X-100 in PBS for 5 minutes at RT, and blocked with 3% BSA in PBST for 30 minutes. Incubation with primary antibody anti-MDM2 (sc-965, 1:500) was done overnight at 4°C. Cell nuclei were counterstained with DAPI. Images were acquired with the High-Content Screening System (ImageXpress) and analyzed with the MetaXpress High-Content Image Analysis Software (Molecular Devices).

### Lentivirus synthesis

HEK293T cells were used for the synthesis of a 3rd-generation lentiviral particles. Briefly, 6×10^6^ cells were transfected in DMEM containing 2mM glutamine and 10% FBS without penicillin or streptomycin. Lipofectamine 2000 was used to transfect 10□μg of the plasmid of interest and the plasmids coding for the lentivirus packaging components (6.5□µg pMMDLRRE, 2.5□µg PRSVREV, and 3.5□µg PMDGVSVG) (#12251, #12253, and #12259, respectively, Addgene). 48 h post-transfection, the viruses were filtered (0.25 µm) and collected.

### Reconstitution of HCT116 *TP53* KO cells with mutant p53

A total of 2×10^5^ HCT116 *TP53* KO cells were seeded in 6-well plates and infected with lentivirus and 10□µg/ml of polybrene, and were incubated for 24□h before changing the media. 2 days after transduction, cells were selected with 10□µg/mL of blasticidin S HCl (A1113903, Gibco) for 10 days. Lentiviral particles were generated with vectors encoding human p53 with a C-terminal V5 tag coding for WT p35, p53R175H, p53R249S, p53R273H, p53R280K (#22945, #22936, #22935, #22934 and #22933, respectively, Addgene, (Junk et al. 2008)) and the control vector pLenti CMV Blast DEST (706-1) (#17451, Addgene).

### Generation of SKOV3 MDM2-overexpressing cells

Cells containing an empty vector (E.V.) or sgRNAs for MDM2 overexpression were generated by using CRISPRa/dCas9-SAM system, which consists of a dead Cas9 (dCas9) fused to the transcriptional activator VP64. First, a total of 2×10^5^ of SKOV3 cells were seeded in 6-well plates and were infected with lentiviral particles containing the lenti dCAS-VP64_Blast plasmid (Addgene #61425). Cells were then selected for ten days with 10 µg/mL of blasticidin. Next, these cells were infected with the pXPR_502 plasmid (Addgene #96923), containing the transcription factors MS2, p65 and HSF1, and a sgRNA targeting the *MDM2* promoter.

Pools of cells were generated by either infecting with an empty vector (E.V.), with no sgRNAs, or with two different sgRNAs directed to different MDM2 promoter regions: MDM2 #1) 5’-GTTCACACTAGTGACCCGAC-3’ and MDM2 #2) 5’-CCCCCACTCCATCATCCCGG-3’. Infected cells were selected with 1 µg/mL puromycin for 7 days.

## Acknowledgements

We thank Dr. Cristina Mayor for providing advice and sharing reagents. This work was supported by grants from the Scientific Foundation of the Spanish Association Against Cancer (PRYGN259366LOPE), the Spanish Ministry of Science and Innovation-Agencia Estatal de Investigacion (PID2020-119329RB-I00 and PID2024-161099OB-I00). Dr. Ivan V Rosadós lab was supported by PID2021-128988OB-I00, CNS2022-136055 and PID2024-161661NB-I00. L.S.-C. was supported with a postdoctoral grant from the Scientific Foundation of the Spanish Association Against Cancer (POSTD211274SIMO). M.L.-P. was supported by the FPU22-00250 and S.R. by the FPU23-00552 fellowships from the Spanish Ministry of Science, Innovation and Universities. E.P. was supported by the by the HORIZON-MSCA ITN RepliFate (101072903).

## Supplementary Figure Legends

**Supplementary Figure 1. PARP1 degradation by a VHL-recruiting PROTAC alleviates Talazoparib-induced cell toxicity.** A) Structure of 180055 PROTAC formed by Rucaparib joined to a VHL ligand. B) Expression of PARP1 in isogenic *VHL* WT and KO KBM7 cells treated with 1µM 180055 for 48h assessed by western blot. Actin was used as a loading control. C) Viability assay by MTT performed in *VHL* WT or KO KBM7 cells pre-treated with 1 µM 180055 for 24 h followed by a treatment with both 180055 and the indicated doses of Talazoparib for another 24 h. Data are normalized to vehicle or PROTAC pre-treated cells and correspond to three biological replicates. IC50s were calculated with non-linear regression analysis and are shown in the figure (µM). Statistical significance was determined with Extra sum-of-squares F test. *** = *p* < 0.001.

**Supplementary Figure 2. Simultaneous treatment with SK-575 and PARPi impacts PROTAC-driven protection.** A) Viability assay by MTT performed with *CRBN* WT or KO KBM7 cells treated simultaneously with 100 nM SK-575 or vehicle, and the indicated doses of Olaparib or Talazoparib for 24h. Data are normalized to vehicle or SK-575 treated cells and correspond to three biological replicates. IC50s were calculated with non-linear regression analysis and are shown in the figure (µM). Statistical significance was determined with Extra sum-of-squares F test. B) Colony-forming cells (CFC) assay of human bone marrow mononuclear cells simultaneously treated with vehicle or 100 nM SK-575 at the time of seeding them in MethoCult matrix with the indicated doses of Talazoparib. Colonies were quantified after 14 days from three replicates. Statistical significance was determined with unpaired t-tests. * = *p* < 0.05 and ** = *p* < 0.01.

**Supplementary Figure 3. Reconstitution of HCT116 *TP53* KO cells with mutant p53 does not rescue BRD4 degradation by A1874 PROTAC.** A) Expression of p53 in cell lines transduced with pLenti6 vector expressing either wildtype or mutant p53 with a V5 tag. B) Expression of BRD4, MDM2, p53 and p21 in HCT116 cells with the indicated genotypes, treated with vehicle or 1 µM A1874 PROTAC for 24 h. Vinculin was used as a loading control.

**Supplementary Figure 4. *TP53*-mutant cancers display reduced *MDM2* mRNA levels.** A) Xena Browser Visual Spreadsheet for TCGA Pan-Cancer (PANCAN), TCGA Colon and Rectal Cancer (COADREAD), TCGA Breast Cancer (BRCA), TCGA Melanoma (SKCM) and TCGA Ovarian Cancer (OV) studies. The left column shows *MDM2* gene expression and the right column shows *TP53* somatic mutations. Expression is colored red to blue for high to low expression. The gene diagram at the top of *MDM2* column shows *MDM2* exons as boxes, with tall coding regions and shorter untranslated regions. The gene diagram of top of *TP53* column shows *TP53* exons as boxes. The position of each mutation is marked in relation to the gene diagram and colored by its functional impact. Null data is not shown (except for the Ovarian cancer cohort, where only samples with null data for both features have been removed). B) Violin plot showing *MDM2* expression level in the indicated TCGA datasets comparing *TP53* wildtype and *TP53* mutant samples. Statistical significance was determined with unpaired t-tests. **** = *p* < 0.0001.

