## Supplementary figures for "PROTAC-Driven Protective Therapy increases the therapeutic window of anticancer drugs"

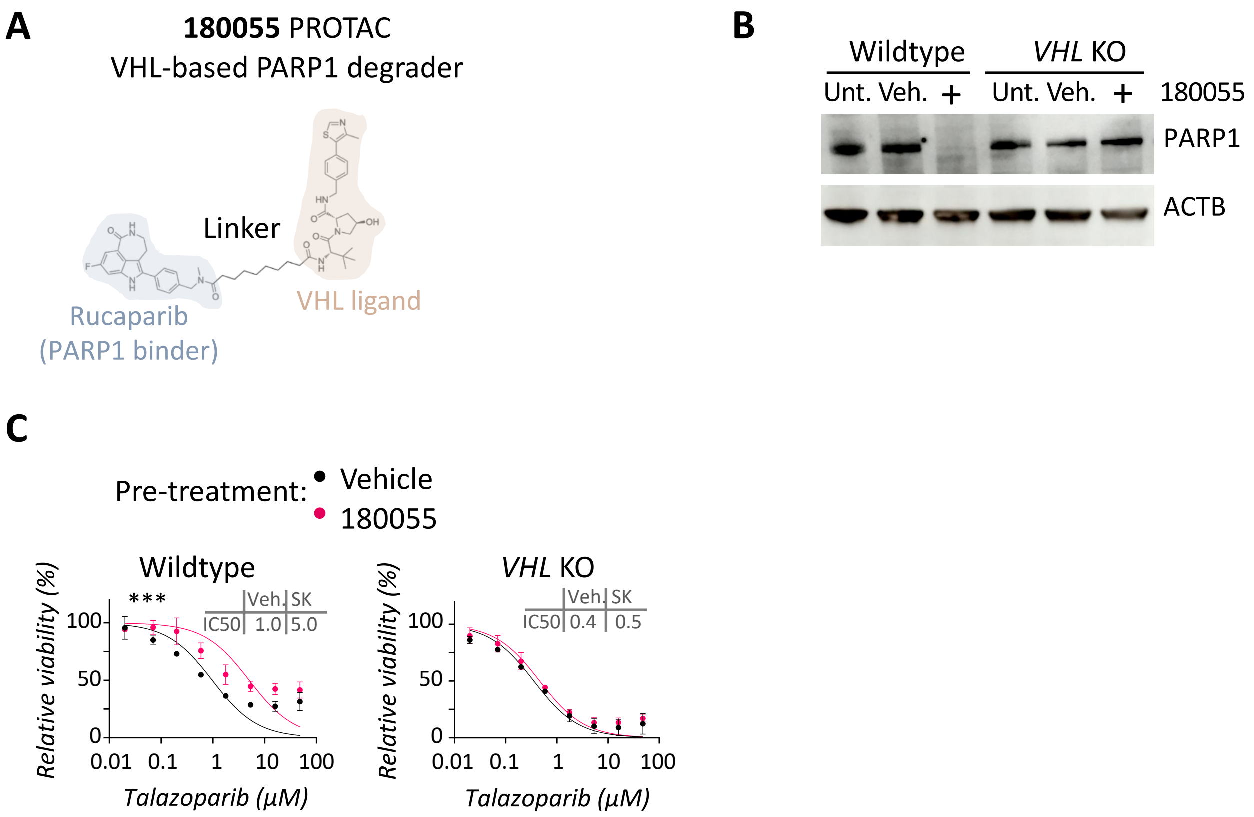


Supplementary Figure 1 - Simón-Carrasco et al.

**Supplementary Figure 1. PARP1 degradation by a VHL-recruiting PROTAC alleviates Talazoparib-induced cell toxicity.** A) Structure of 180055 PROTAC formed by Rucaparib and a VHL ligand. B) Expression of PARP1 in isogenic *VHL* WT and KO KBM7 cells treated with 1µM 180055 for 48h assessed by western blot. Actin was used as a loading control. C) Viability assay by MTT performed in *VHL* WT or KO KBM7 cells pre-treated with 1 µM 180055 for 24 h followed by a treatment with both 180055 and the indicated doses of Talazoparib for another 24 h. Data are normalized to vehicle or PROTAC pre-treated cells and correspond to three biological replicates. IC50s were calculated with non-linear regression analysis and are shown in the figure (µM). Statistical significance was determined with Extra sum-of-squares F test. *** = $p < 0.001$.

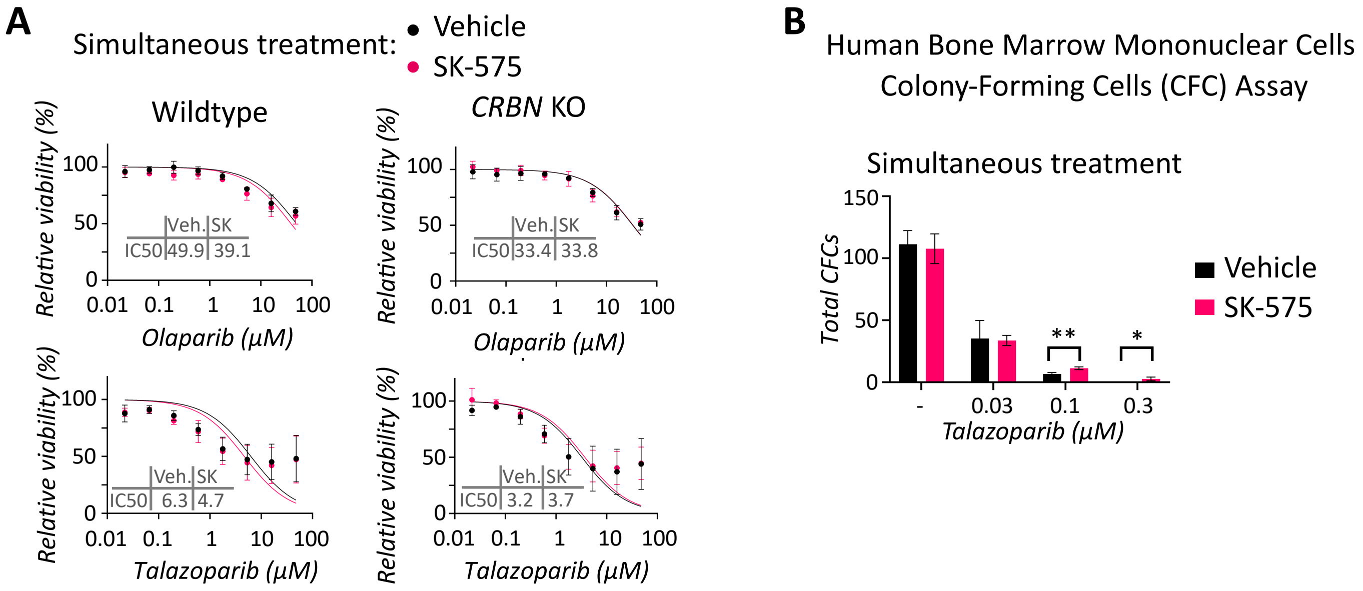


**Supplementary Figure 2 - Simón-Carrasco et al.**

**Supplementary Figure 2. Simultaneous treatment with SK-575 and PARPi impacts PROTAC-driven protection.** A) Viability assay by MTT performed with *CRBN* WT or KO KBM7 cells treated simultaneously with 100 nM SK-575 or vehicle, and the indicated doses of Olaparib or Talazoparib for 24h. Data are normalized to vehicle or SK-575 treated cells and correspond to three biological replicates. IC50s were calculated with non-linear regression analysis and are shown in the figure (µM). Statistical significance was determined with Extra sum-of-squares F test. B) Colony-forming cells (CFC) assay of human bone marrow mononuclear cells simultaneously treated with vehicle or 100 nM SK-575 at the time of seeding them in MethoCult matrix with the indicated doses of Talazoparib. Colonies were quantified after 14 days from three replicates. Statistical significance was determined with unpaired t-tests. * = $p < 0.05$ and ** = $p < 0.01$.

A

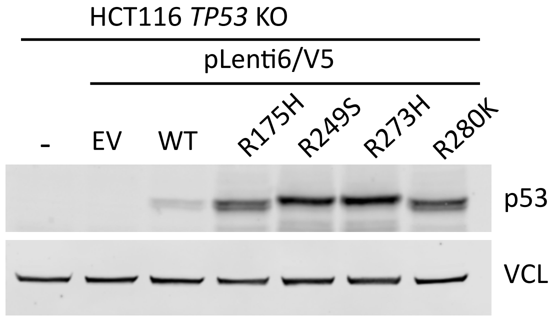


B

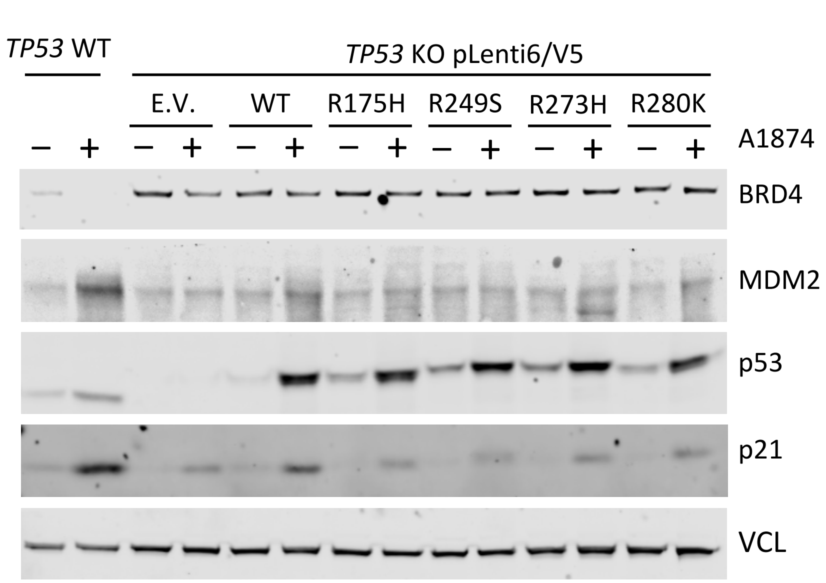


**Supplementary Figure 3 - Simón-Carrasco et al.**

**Supplementary Figure 3. Reconstitution of HCT116 *TP53* KO cells with mutant p53 does not rescue BRD4 degradation by A1874 PROTAC.** A) Expression of p53 in cell lines transduced with pLenti6 vector expressing either wildtype or mutant p53 with a V5 tag. B) Expression of BRD4, MDM2, p53 and p21 in HCT116 cells with the indicated genotypes, treated with vehicle or 1 μM A1874 PROTAC for 24 h. Vinculin was used as a loading control.

**A**

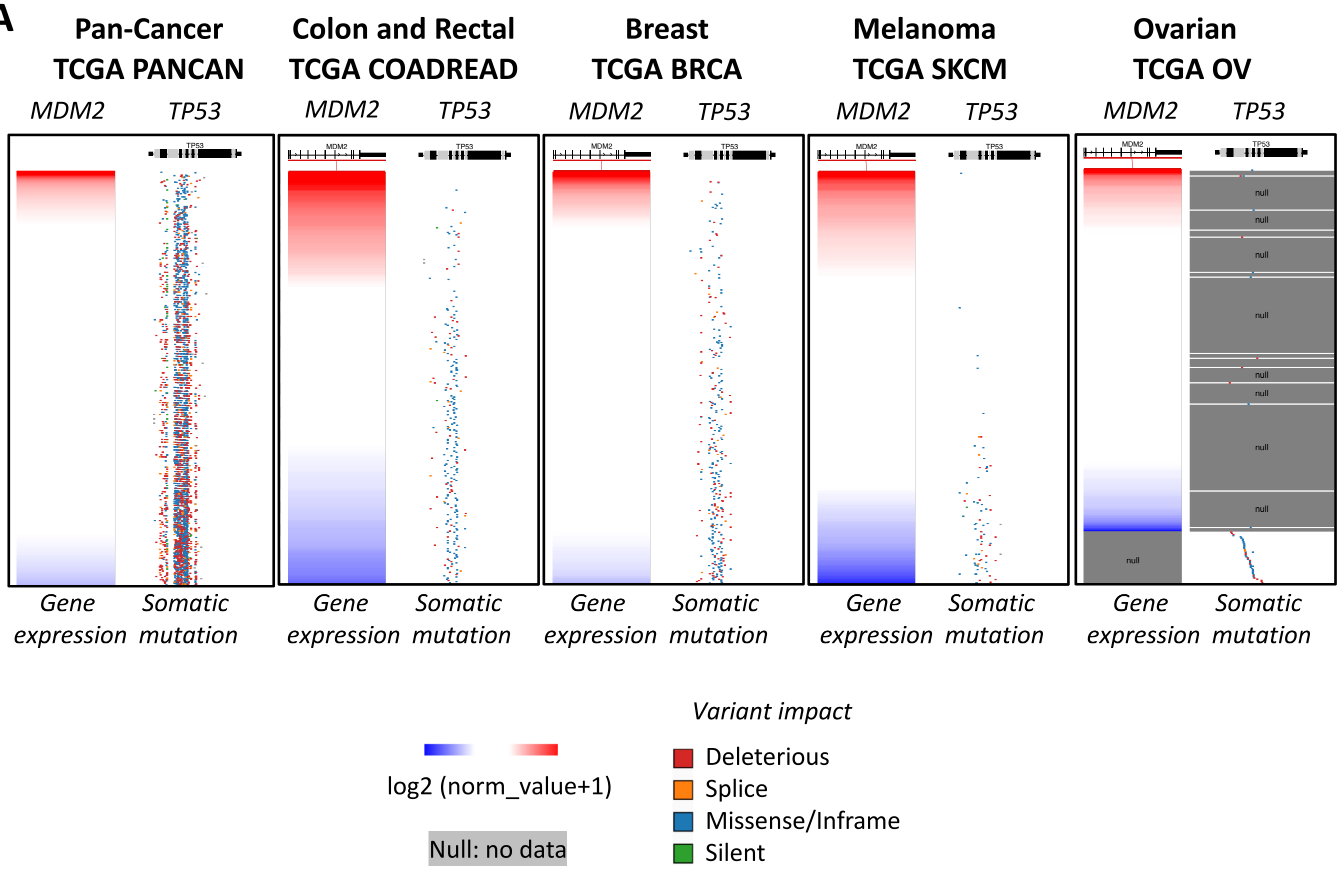


**B**

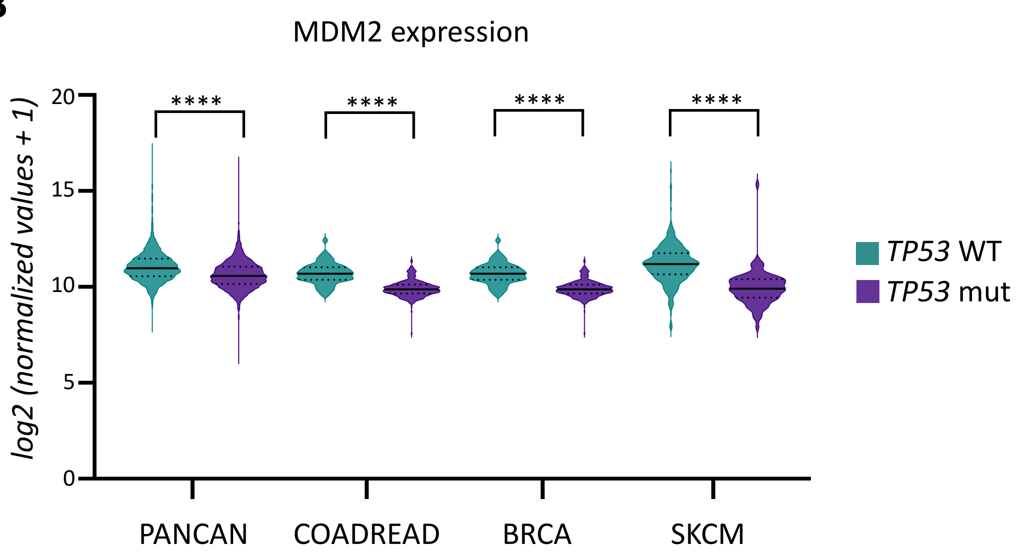


**Supplementary Figure 4 - Simón-Carrasco et al.**

**Supplementary Figure 4**. ***TP53*-mutant cancers display reduced *MDM2* mRNA levels.** A) Xena Browser Visual Spreadsheet for TCGA Pan-Cancer (PANCAN), TCGA Colon and Rectal Cancer (COADREAD), TCGA Breast Cancer (BRCA), TCGA Melanoma (SKCM) and TCGA Ovarian Cancer (OV) studies. The left column shows *MDM2* gene expression and the right column shows *TP53* somatic mutations. Expression is colored red to blue for high to low expression. The gene diagram at the top of *MDM2* column shows *MDM2* exons as boxes, with tall coding regions and shorter untranslated regions. The gene diagram of top of *TP53* column shows *TP53* exons as boxes. The position of each mutation is marked in relation to the gene diagram and colored by its functional impact. Null data is not shown (except for the Ovarian cancer cohort, where only samples with null data for both features have been removed). B) Violin plot showing *MDM2* expression level in the indicated TCGA datasets comparing *TP53* wildtype and *TP53* mutant samples. Statistical significance was determined with unpaired t-tests. **** = $p < 0.0001$.
